# Weight loss response following lifestyle intervention associated with baseline gut metagenomic signature in humans

**DOI:** 10.1101/2021.01.05.425474

**Authors:** Christian Diener, Shizhen Qin, Yong Zhou, Sushmita Patwardhan, Li Tang, Jennifer Lovejoy, Andrew T. Magis, Nathan D. Price, Leroy Hood, Sean M. Gibbons

## Abstract

We report a weight-loss response analysis on a small cohort of individuals (N=25) selected from a larger population (N∼5,000) enrolled in a commercial scientific wellness program, which included healthy lifestyle coaching. Each individual had baseline data on blood metabolomics, blood proteomics, clinical labs, lifestyle questionnaires, and stool metagenomes. A subset of these participants (N=15) lost at least 10% of their body weight within a 6-12 month period and saw significant improvement in metabolic health markers (‘weight loss’ group), while another subset of individuals (N=10) undergoing the same lifestyle intervention showed no change in BMI over the same timeframe (‘no weight loss’ group). Only a single baseline blood analyte, a metabolite linked to fried food consumption, was (negatively) associated with weight loss, but a large number of baseline stool metagenomic features, including complex polysaccharide and protein degradation genes, stress-response genes, respiration-related genes, cell wall synthesis genes, and gut bacterial replication rates, were significantly associated with weight loss after explicitly controlling for baseline BMI. Together, these results provide a set of baseline gut microbiome functional features that are associated with weight loss outcomes.

## Main Text

The question of whether or not there are consistent associations between the ecological composition of the gut microbiome and human obesity remains somewhat controversial ^1^. There are many confounding variables with regard to obesity phenotypes, including genetics, prior health status, age, physical activity, and diet, which can modulate whether or not a person who is nominally ‘overweight’ or ‘obese’ is considered metabolically healthy ^2–4^. Assuming an association between the microbiota and obesity exists, another important, but unresolved, question is whether or not the human gut microbiome is simply diagnostic of obesity or if our microbiota somehow contribute directly to host weight change. While the gut microbiome has been shown to contribute to weight gain in mice ^5^, its potential contribution to weight gain/loss in humans is still poorly understood ^6^. Recent feeding studies have shed some light on this issue, demonstrating that humans with higher *Prevotella*-to-*Bacteroides* ratios tend to lose significantly more weight on a high-fiber diet ^7^. Thus, while the mechanisms remain unclear, the baseline composition of the human gut microbiota can indeed influence host responses to weight-loss interventions. Finally, it is unclear if the associations between the gut microbiome and weight loss phenotypes are independent of the effects of baseline BMI on the microbiome. In this pilot study, we set out to better understand the potential interactions between BMI, metabolic health, weight loss, and gut microbiome functional profiles in data from a human cohort that underwent a healthy lifestyle intervention.

For this study, we leveraged existing data and biobanked samples from the Arivale cohort (see Methods). Briefly, participants enrolled in a commercial behavioral coaching program run by the former scientific wellness company Arivale, Inc., were paired with a registered dietitian or registered nurse coach. Personalized, telephonic behavioral coaching was provided to each participant on a monthly basis, with email or text communications between coaching calls. This service included longitudinal ‘deep phenotyping’, which involved collecting blood and stool for baseline SNP genotyping or whole genome sequencing (blood) and longitudinal clinical labs (blood), metabolomics (blood), proteomics (blood), and 16S amplicon sequencing of the microbiome (stool), along with lifestyle questionnaires, body weight measurements, and additional activity-tracking data from wearable devices. Arivale participants showed broad improvements across a number of validated health markers while enrolled in the program, including an average reduction in BMI ^8,9^.

We selected a subset of the ∼5,000 Arivale participants to look specifically at weight loss phenotypes during the lifestyle intervention period (Fig. 1A). Briefly, we examined 1,252 individuals with blood collected at two timepoints over the course of a year, 239 of which had a paired stool sample at baseline and longitudinal data on BMI (Fig. 1A-C). We further subdivided these 239 participants by selecting individuals who lost > 1% of their body weight per month over a 6-12 month period (N=15) and those who maintained a stable BMI (N=10) over the same period (Fig. 1C). Biobanked fecal samples from this 25 person cohort were used to generate shallow shotgun metagenomes, in order to obtain gut microbiome functional and taxonomic profiles. Biobanked plasma samples from these same individuals were used to generate additional proteomic data on a broad set of obesity and cardiometabolic health markers (Table S1). In the ‘weight loss’ group, participants lost an average of 1.5-4.5% of their body weight per month over 10-12 months (>10% overall), while the ‘no weight loss’ group showed no significant change in weight during the intervention (Fig. 1D). There was no significant difference in age between ‘weight loss’ and ‘no weight loss’ groups, but the ‘weight loss’ group had a significantly higher baseline BMI (Fig. 1E-F). All individuals in the ‘weight loss’ group were considered either overweight or obese (BMI > 25 and 30, respectively), while half of the ‘no weight loss’ group were overweight and the other half were considered normal weight (BMI > 25 and < 25, respectively; Fig. 1E). Despite differences between groups in baseline BMI, baseline markers of cardiometabolic health, like serum LDL cholesterol, serum adiponectin, and serum glucose levels, were not significantly different between groups (Fig. 1G-I).

**Figure 1.**
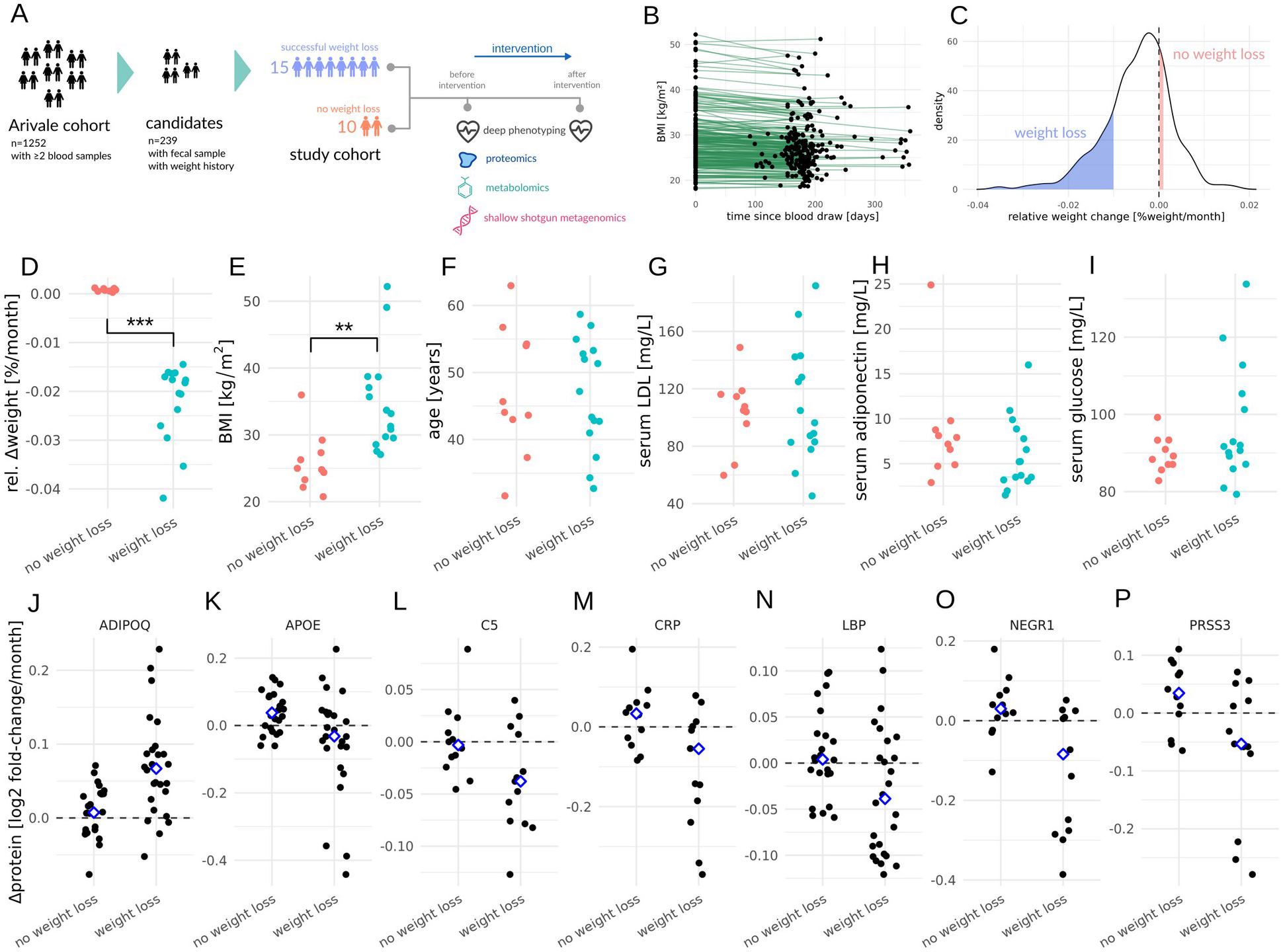
Cohort selection for weight loss analysis. Schematic showing number of individuals within the Arivale wellness intervention cohort that match our selection criteria for ‘weight loss’ and ‘no weight loss’ groups (A). Baseline and follow-up BMI values (points from same individual connected by lines) for 239 individuals with the full set of ‘omics and biometric data available (B). Distribution of relative weight change for the same 239 individuals, with blue area showing individuals who lost >1% of their body weight per month (N=15) and red area showing individuals who showed not change in weight (N=10) over the same intervention period (C). Dot plots showing relative weight changes (D), baseline BMI (E), age (F), baseline serum LDL (G), baseline serum adiponectin (H), and baseline serum glucose (I) in the ‘weight loss’ and ‘no weight loss’ groups. Blood proteins with significantly different slopes (FDR-corrected p<0.1) between baseline and follow up sampling in the ‘weight loss’ group (J-P). Dashed line denotes no change in protein abundance over time and blue diamonds denote the means of the two groups (J-P).

On average, only individuals in the ‘weight loss’ group showed broad improvements in blood proteomic markers of obesity and cardiometabolic health (FDR-corrected ANOVA p<0.1, Fig. 1J-P). Specifically, the weight loss group showed a marked increase in ADIPOQ (adiponectin) levels, which have previously been negatively associated with BMI and positively associated with fasting ^10^. The weight loss group also showed decreased levels of APOE, C5, CRP, NEGR1, and PRSS3, which have all been positively associated with obesity, inflammation, and metabolic disorders (Fig. 1J-P) ^10–14^. Thus, not only did the weight loss group reduce their BMI during the intervention period, but they became metabolically and immunologically healthier as well.

We tested for associations between baseline features and weight loss that were independent of BMI, age, and sex (Fig. 2A). Even though one might expect baseline factors associated with BMI to have a similar association with weight loss (see Fig. 2A), we found that these two effects were largely independent for baseline metabolomics and proteomics features (Pearson rho ∼ 0.1) and only weakly correlated for baseline gut microbial species abundances and genes (Pearson rho = 0.22 and 0.3, respectively), indicating that baseline ‘omics feature associations with BMI and weight loss are mostly orthogonal (Fig. 2B-E). Few baseline blood metabolomic features were significantly associated with BMI, and only one unclassified metabolite (Metabolon ID: X-11378) was independently associated with weight loss (Fig. 2B). Interestingly, X-11378 has previously been associated with the consumption of fried foods ^15^, and was negatively associated with weight loss in our study. Only one baseline blood protein feature was negatively associated with BMI, and no proteins were independently associated with weight loss (Fig. 2C). In contrast to blood-derived features, many gut bacterial taxa were significantly associated with BMI, although no taxa were independently associated with weight loss (Fig. 2D). However, many gut bacterial functional genes showed independent associations with either BMI or weight loss, and a few showed independent associations with both BMI and weight loss (Fig. 2E). Prior work in a larger cohort of several hundred individuals showed how blood analytes could predict glycemic responders during a clinical weight loss program ^4^. However, that study did not look at the baseline gut microbiomes. Here, we find that the baseline stool metagenome shows a much stronger association with weight loss phenotypes than baseline blood proteomic or metabolomic features (Fig. 2B-E).

**Figure 2.**
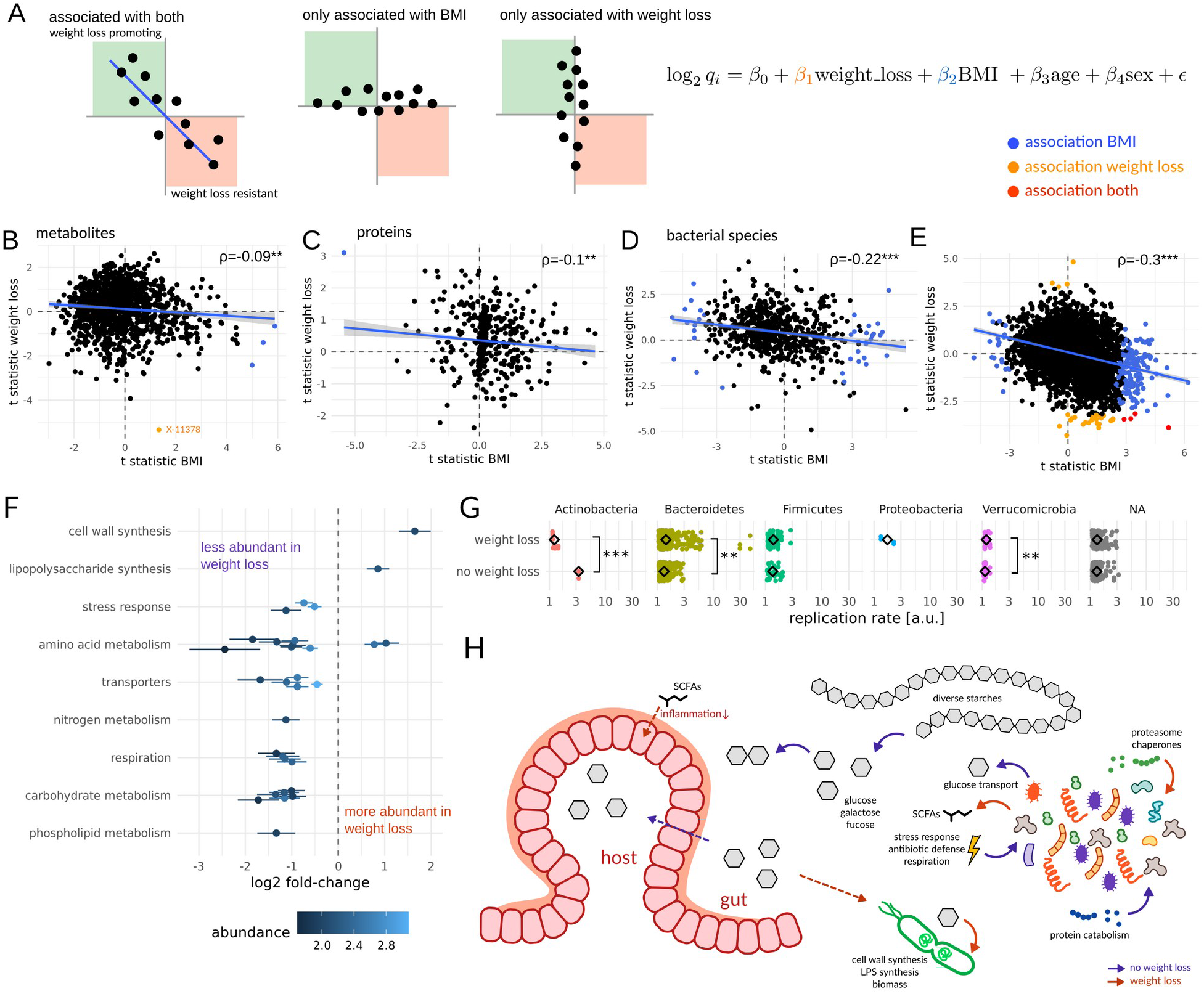
Associations between baseline multi-omic features, BMI, and weight loss. Biplots show t-statistics for features associated with BMI or weight loss, controlling for age and sex (A). Analyses were run separately for blood metabolites (B), blood proteins (C), gut bacterial taxa (D), and gut bacterial functional genes (E). Blue dots denote features significantly associated with BMI only (i.e. independent of weight loss, age, and sex), orange dots denote features significantly associated with weight loss only (independent of BMI, age, and sex), and red dots denote features independently associated with both BMI and weight loss. Metagenomic features significantly associated with weight loss, binned into high-level functional categories (F). Average phylum-specific bacterial replication rates estimated from metagenomes show significant differences across weight loss groups (G). Schematic of specific metagenomic functions from panel F that were positively or negatively associated with weight loss and how they might be involved in modulating dietary energy buffering or gut inflammation (H).

Thirty-two baseline gut microbiome functional genes were independently associated with weight loss (Fig. 2F). Cell wall and lipopolysaccharide (LPS) synthesis were positively associated with weight loss, which suggested that cell division, biomass production, and gram negative bacterial growth potential might be important. To explore this further, we calculated baseline bacterial replication rates directly from metagenomically-assembled contigs ^16^, and found that average replication rates were indeed significantly higher in the ‘weight loss’ group, with gram negative Bacteroidetes contigs contributing most to this effect (Fig. 2G). Most contigs could not be annotated beyond the phylum level, but the fastest replicating contigs (replication rate > 3) with genus-level annotations belonged to *Prevotella*, and were enriched in the weight-loss group. Most functional genes were negatively associated with weight loss: specifically, functions involved in glycan and protein catabolism, response to stress, peptide antibiotic synthesis, and respiration (Fig. 2H).

Based on our results, we propose a tentative model whereby gut commensals modulate the host’s absorption of calories from the diet and potentially impact intestinal inflammation (Fig. 2H). Specifically, we know that the gut microbiota help break down complex, extracellular polysaccharides into simpler sugars that are more readily absorbed by the host.

If commensal gut bacterial growth rates are reduced, the host epithelium may be able to better compete with commensals for these extracellular breakdown products before they can be transformed into less-energy-dense fermentation byproducts, like short-chain-fatty-acids (SCFAs), and bacterial biomass. SCFA production itself can reduce intestinal inflammation ^17^, which in turn may help to improve metabolic health and better facilitate weight loss ^18^.

Concordantly, we saw reduced levels of circulating inflammation-related proteins in participants who lost weight (Fig. 1L-M). Finally, reduced inflammation could itself promote fermentative metabolism and redox homeostasis in the gut, minimizing oxygen stress to strict anaerobes, and suppressing respiratory pathways that favor facultative anaerobes (Fig. 2H).

In summary, we suggest that dietary energy buffering, host-microbe substrate competition, and modulation of host inflammation by commensal bacteria may be, in part, responsible for determining host responses to healthy lifestyle interventions. Gut ecosystems optimized for fermentative metabolism and higher bacterial growth rates appear to be conducive to weight loss. Prior work has shown that the higher baseline levels of *Prevotella* can improve weight loss responses to a standardized high-fiber diet ^7^, and here we found higher baseline *Prevotella* growth rates in individuals who lost weight in a non-standardized wellness program, which often involved suggested increases in dietary fiber. Perhaps unsurprisingly, we saw that a metabolite associated with eating fried foods was negatively associated with weight loss.

Ultimately, we will need more data from human interventional trials to improve our understanding of how our commensal gut microbiota and our diet causally contribute to weight loss. By combining these emerging insights with recently developed models for predicting personalized gut microbiome metabolic outputs ^19,20^, we can begin to engineer the functional capacity of our microbiota to optimize dietary and lifestyle interventions.

## Methods

### Arivale cohort and sub-cohort selection criteria

Procedures for this study were run under the Western Institutional Review Board (WIRB) with Institutional Review Board (IRB) study number 20170658 at the Institute for Systems Biology and 1178906 at Arivale. The research was performed entirely using de-identified and aggregated data of individuals who had signed a research authorization allowing the use of their anonymized data in research. Per current U.S. regulations for use of deidentified data, informed consent was not required. To be eligible to join the program, participants had to be over 18 years of age, not pregnant, and a resident of any U.S. state except New York. The participants analyzed in this study are the 92% of participants who agreed to research use as of 6/19/2018 and enrolled in the program between July 2015 and March 2018.

Of the ∼5,000 Arivale participants who agreed to research use of their data, 1,252 had blood draws at two time points (i.e. a baseline sample and then a follow-up sample at ∼6-12 months). Of these 1,252 individuals, 239 had follow-up BMI data within the year after the first blood draw and had biobanked serum and fecal samples available which were samples within 30 days of each other. From those we removed individuals with zero variance in weight measurements, which results from digital scales when regular weighing is not performed and the prior weight is reported repeatedly. Relative weight change was calculated as [follow up weight - baseline weight] / months between measurements. The 15 individuals with the largest drops in weight were used as the “weight loss” group, whereas 10 individuals with the 20 smallest positive weight change values were chosen as the “no weight loss” group (10-person subset was selected to ensure a balanced representation of sexes across groups).

### Arivale behavioral intervention

Participants who enrolled in the year-long commercial behavioral coaching program were paired with a registered dietitian or registered nurse coach. Personalized, telephonic behavioral coaching was provided to each participant on a monthly basis, with email or text communications between coaching calls. Each participant’s clinical and genetic data were available to them via a web dashboard and mobile app, which they could also use to communicate with their coach and schedule calls or blood draws. Coaches provided specific recommendations to address out-of-range clinical results based on clinical practice guidelines, published scientific evidence, or professional society guidelines. Examples of the evidence behind the coaching recommendations include guidelines from the American Heart Association or American Diabetes Association ^21^, comprehensive lifestyle interventions such as those developed for the Diabetes Prevention Program (DPP) ^22^, nutrition recommendations such as those based on the DASH dietary pattern ^23^ or MIND Diet ^24^, and exercise recommendations from the American College of Sports Medicine ^25^.

### Blood collection and multi-omic data generation

Blood draws for all assays were performed by trained phlebotomists at LabCorp or Quest service centers and were scheduled every 6 months, but actual collection times varied. Metabolon conducted the metabolomics assays on participant plasma samples. Sample handling, quality control and data extraction, along with biochemical identification, data curation, quantification and data normalizations have been previously described ^26^. For analysis, the raw metabolomics data were median scaled within each batch, such that the median value for each metabolite was one. To adjust for possible batch effects, further normalization across batches was performed by dividing the median-scaled value of each metabolite by the corresponding average value for the same metabolite in quality control samples of the same batch. Missing values for metabolites were imputed to be the minimum observed value for that metabolite. Values for each metabolite were log transformed. Plasma protein levels were measured using the ProSeek Cardiovascular II, Cardiovascular III and Inflammation arrays (Olink Biosciences), processed and batch corrected as described previously ^26^. For analysis, a threshold of less than 5% missing values was set for each protein, which was passed by 263 different analytes. Missing values for the proteins were imputed to be the minimum observed value for that protein.

### Stool collection and metagenomic data generation

At home stool collection kits (DNAgenotek, OMR-200) were shipped directly to participants, and then shipped back to DNA Genotek for processing. Microbial DNA was isolated from 200 μL of homogenized fecal material using the DNeasy PowerSoil Pro extraction kit (Qiagen, Germany) with bead beating in Qiagen Powerbead Pro plates (Cat No. 19311, Qiagen, Germany). Extracted DNA was quantified using the Quant-iT PicoGreen dsDNA Assay kit (Invitrogen, USA) and all samples passed the quality threshold of 1ng/μL (range 8-101ng/μL). Shallow shotgun sequencing was performed with the BoosterShot service (Corebiome, USA). In brief, single-stranded 100bp libraries were prepared using an optimized proprietary protocol of the provider (Corebiome, USA) based on the Nextera library prep kit (Illumina, USA) and sequenced on a NovaSeq (Illumina, USA) to a minimum of 2M reads per sample. Demultiplexing was performed on Basespace (Illumina, USA) yielding the final FASTQ files.

### Anthropometric Data

Height, weight, and waist circumference were measured either at the blood draws (45%), or were self-reported via an online assessment, or through the Fitbit Aria scale. Reference ranges for anthropometric data were defined by U.S. public health guidelines ^27^.

### Selective Reaction Monitoring (SRM) of obesity-related proteins

Serum samples were processed following a previously published protocol that ensured maximum yield of signal ^28^. We targeted a curated selection of 22 mostly organ-specific proteins with known genetic variants associated with obesity or metabolic syndrome (Table S1). Prepared samples, along with spiked-in heavy isotope labeled synthetic standard peptides, were quantified using a triple-quadrupole mass spectrometer (Agilent 6490, Agilent, Santa Clara, CA) with a nanospray ion source and Chip Cube nano-HPLC. Three to four transitions were monitored for each target peptide (see Table S1). Two ug of tryptic digested Mars-14 (Agilent, Santa Clara, CA) depleted serum were eluted from a high-capacity nano-HPLC Chip (160 nL, 150 mm × 75 μm ID, Agilent, Santa Clara, CA) with a 30 min gradient of 3-40% acetonitrile as described previously ^28,29^. Raw SRM mass spectrometry data was analyzed with the Skyline targeted proteomics environment ^30^. Each detected peptide was quantified by the light/heavy (L/H) ratio of monitored transitions, after adjusting for the volume of the original serum sample.

### Metagenomics data processing

Trimming and filtering for the raw sequencing data was performed using FASTP v0.20.1 ^31^. The first five bases on the 5’ were trimmed from each read to avoid leftover PCR primers and each read was furthermore trimmed on the 3’ by sliding window method with a minimum quality threshold of 20. Abundances of species were obtained using KRAKEN v2.0.9 and BRACKEN v2.6.0 using the default KRAKEN database ^32,33^. Contigs were assembled *de novo* with MEGAHIT v1.2.9 with a cross-assembly across all samples. Open reading frames (ORFs) in the resulting contigs were then identified with PRODIGAL v2.6.3 ^34^. Reads from each sample were then aligned to each contig using MINIMAP2 v2.17 and gene abundances for each sample were quantified with the Expectation-Maximization algorithm from SALMON v3.1.3 ^35,36^. The identified ORFs were annotated using the EGGNOG EMAPPER v2.0.1.

Replication rates were inferred using the iRep approach ^16^. Here, we first aligned the reads for each sample to the full assembled contigs using MINIMAP2 v2.17. Coverage profiles were extracted for all contigs larger than 5000bps across bins of a 100bp width but only contigs with a minimum length of 11000bp and a mean coverage of 2x were used for the iRep inference. Coverage profiles were smoothened using a sliding window mean over 50 bins (5000bp window width) before calculating the replication rates using the iRep implementation in mbtools v0.44.14 (https://gibbons-lab.github.io/mbtools). Taxonomic classification of individual contigs were obtained using CAT v5.1.2 using the included database of single-copy marker genes ^37^.

### Statistical analyses

#### SRM data

Raw SRM abundances were log-transformed and as this yielded data that appeared to be normal distributed (as validated by QQ-plots). Change in protein abundance across the intervention was then quantified as the difference of protein abundance after intervention and the baseline abundance, yielding log ratios of post-intervention vs. baseline abundances.

Associations with weight loss were obtained by linear regression of the obtained log ratios using the design shown in Fig. 2. Here, assignment to weight loss group was the target covariate corrected for baseline BMI, age and sex. False discovery rates were controlled by adjusting p-values using the Benjamini-Hochberg correction.

#### Metabolomic and proteomic data

Mass spectroscopy data from untargeted metabolomics and proteomics was log-transformed as this yielded near-normal distributions on QQ plots. Log-abundance values were then used for linear regressions using the design formula shown in Fig. 2. For each metabolite and protein we also performed a regression without the weight loss group and using the baseline BMI as the target covariate to yield the association strength with BMI. Linear regressions were run using the LIMMA R package without Bayesian shrinkage as this is specific to gene expression data ^38^. False discovery rates were controlled by adjusting p-values using the Benjamini-Hochberg correction. T-values for each association coefficient were calculated as the ratio of coefficient and estimated coefficient standard deviation obtained from the Fisher matrix of the regression.

#### Metagenomic data

As species abundances as well as gene abundances were both obtained from sequencing count data we analyzed both data types using negative binomial regressions which have been shown to fit metagenomic data well ^39^. This again used the design shown in Fig. 2. However, this time the regressions were performed with negative binomial regression using DESeq2 and using a prior normalization (“poscounts” method in DESeq2) ^40^. For each metagenomic feature (species or gene) we also performed a regression without the weight loss group and using the baseline BMI as the target covariate to yield the association strength with BMI. Pseudo t-values were calculated as the ratio of coefficient and estimated coefficient standard deviation obtained from DESeq2.

### Data and code availability

Raw metagenomic sequencing data has been deposited on the NCBI Sequence Read Archive (SRA) under Bioproject PROJXXX (currently being uploaded and accession will be provided prior to publication). SRM data can be found on the GitHub repository associated with this study (https://github.com/gibbons-lab/weight_loss_2019). The Institute for Systems Biology manages all Arivale data requests for non-profit research purposes and will grant access to qualified researchers. Data requests should be sent to: A.M.

The full pipeline used to process the metagenomic data is provided as a Nextflow pipeline in the study repository at https://github.com/gibbons-lab/weight_loss_2019. All analyses can be found in Rmarkdown notebooks and allow the reproduction of all analyses and figures in this manuscript. Specialized functions, such as the implementation to calculate replication rates or association analyses, can be found in a dedicated R package along with documentation at https://github.com/gibbons-lab/mbtools.

## Acknowledgements

This work was supported by an Institute for Systems Biology Innovator Award (PIs CD and SQ). SMG and CD were supported by the Washington Research Foundation Distinguished Investigator Award and startup funds from the Institute for Systems Biology.

## Supplemental Information

**Table S1.**
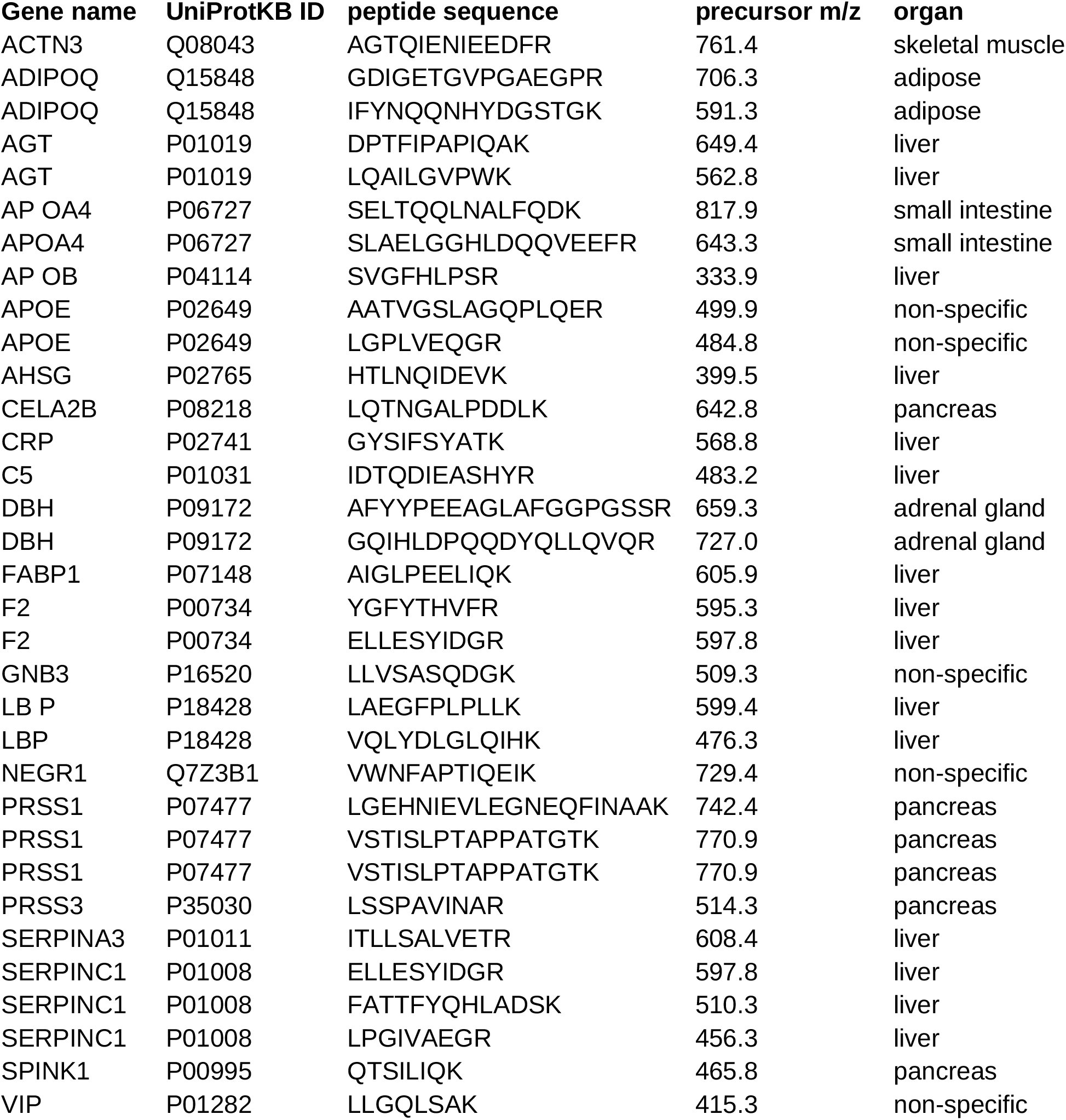
Peptide sequences and transitions for the SRM quantification.

